# Different Perspective of Machine Learning Technique to Better Predict Breast Cancer Survival

**DOI:** 10.1101/2020.07.03.186890

**Authors:** Kaushal Kumar, Vijay Vikram Singh, Ramakrishna Ramaswamy

## Abstract

Machine learning (ML) plays a key job in the guide of cancer diagnosis and identification. The researcher has implemented different algorithms of ML for the prediction of breast cancer. Some researchers recommend their algorithms are more accurate, faster, and easier than others. My study relies on recently developed machine learning algorithms like genetic algorithms and deep belief nets. I’m interested to build a framework to precisely separate among benign and malignant tumors. We’ve optimized the training algorithm. During this unique circumstance, we applied the genetic algorithm procedure to settle on the main genuine highlights and perfect boundary estimations of the AI classifiers. The examinations rely upon affectability, cross-validation, precision, and ROC curve. Among all the varying kinds of classifiers used in this paper genetic programming is the premier viable model for highlight determination and classifier.

## I. Introduction

The study of cancer research isn’t new using Machine learning. Neural networks and decision trees are employed in cancer detection and diagnosis for nearly 20 years. Today the job of AI techniques has used in an exceedingly huge choice of uses beginning from identifying and grouping tumors through X-beam and CRT pictures to the arrangement of malignancies from proteomic and genomic (microarray) measures [1]. In line with the foremost recent PubMed statistics, over 1600 papers are published related to machine learning and cancer [2]. These papers have the enthusiasm to recognize, characterize, identify, or recognize tumors and different malignancies utilizing AI strategies. In other words, Machine learning plays a key job in the guide of cancer diagnosis and identification [3].

Breast cancer could also be a prevalent explanation for death, widespread among women worldwide [4]. Various procedures are produced for early identification and cure of breast cancer growth and diminished the cases of deaths, and different aided breast cancer determination techniques know about increment the diagnostic accuracy [5]. Within the previous number of decades, Machine learning procedures which require three primary stages: preprocessing, classification, and feature extraction. During this process, SVD or PCA is applied to downsize the dimension of the feature vector. Numerous works have endeavored to robotize the determination of breast cancer growth supported AI algorithms [6].

Nowadays, machine learning is playing a very important role in the medical sciences. Unfortunately, AI keeps on being a field with high boundaries and here and there requires master information. Structuring a legit AI model including the phases of preprocessing, highlight determination, and order forms requires a lot of abilities and skill. The master in AI picks the worthy strategy for this difficult space. Notwithstanding, the nonexperts in AI invest numerous some energy to enhance their proposed models and to accomplish the objective execution. During this specific circumstance, the point of the work is to mechanize the structure of the AI models utilizing many procedures.. Genetic programming is the best combination to optimized the techniques. There are two significant worries to mechanize the breast cancer analysis: (i) to find out which model most closely fits the information and (ii) a way to automatically design machine learning model by adjusting the parameters.

In Section 2,3,4 we have explained material methods and mathematical details of algorithms. Section 5 explains the computational results, whereas the foremost conclusions are discussed in Section 6.

## II. mathematical details of algorithms

### Dataset Used

We utilize the breast cancer, Coimbra, Data Set downloaded from the UCI ML Repository. This dataset [7] utilized by Patricio to anticipate harmful and noncancerous tumors. A blood test was totally gathered at the indistinguishable time after for the time being fasting. The clinical, segment, and human-centric information were gathered for all members, under comparable conditions, consistently by the indistinguishable examination doctor during the conference. Gathered information included weight, age, tallness, and menopausal status. During this procedure, clinical highlights, including BMI, age, HOMA, Insulin, Leptin, Resistin, Adiponectin, and MCP-1 were watched or estimated for all of the 166 patients [7]. There are 10 indicators showing the nearness or nonattendance of breast cancer. The indicators are human-centric information and boundaries which may to some degree be assembled in routine blood examination [8].

For analysis purposes, we use several machine learning techniques like Linear SVM, Logistical regression, naive Bayes, random forest, decision tree and KNN [9] to get possible biomarkers.

In this procedure, we attempt to locate a base number of viable features. Feature selection strategies could even be separated into the covering, channel, and installed techniques. During this content, we make the intrigued peruser caution to the probabilities of feature selection [10], giving an essential scientific classification of feature selection strategies, and talking about their utilization, assortment, and, potential during diffusion of both common similarly as upcoming bioinformatics applications [11].

### A. Genetic Algorithm

In 1975 Holland built up the Genetic Algorithm [12]. The Genetic Algorithm (GA) could even be a computational improvement worldview displayed on the idea of biological evolution. GA is a streamlining methodology that works in twofold inquiry spaces and controls a populace of likely arrangements. It assesses fitness function to a get above-average performance with high performance [13].

GA framework encodes the arrangement into a numerical structure considered a chromosome that comprises of the tiniest genetic component of the gene [14]. At that time, haphazardly chose to border a diffusion of chromosomes due to the initial population. At that time, there’ll be chosen sets of chromosomes from the population for crossover and mutation operations [13], rehashed until the result stops inside the perfect condition that the foremost extreme number of generations is accomplished. From that time forward, the fitness values estimations of each chromosome are determined. The preeminent successful fitness value of the chromosome is visiting be chosen inside the gene column for next-generation reproduction [13] [15]. The preeminent successful fitness value of the chromosome is visiting be chosen inside the gene column for next-generation reproduction. The calculations of this technique could even be portrayed as following steps:

1. Generating the beginning population.
2. Evaluate fitness value.
3. Selection process from the underlying populace to lead hybrid operations.
4. Crossover of each pair of chosen parent chromosomes.
5. Mutation on the picked chromosome, and figure the well-ness estimation of the chromosome.
6. After the change, assessing the wellness estimation of the recently produced youngster populace. In the event that the wellness estimation of the posterity is higher, at that point, the parent chromosome is supplanted with a substitution posterity chromosome. On the off chance that not, at that point don’t go spinning.
7. If its bomb rehashes the progression (3) to (6) until it arrives at the preeminent number of ages.

### B. Classification Using Support Vector Machines (SVM)

In 1998 Vapnik built up the SVM. It builds a classifier which partitions preparing tests. And furthermore, boosts the base separation or edge [16]. Think about the arrangement of two classes, which can to some degree be portrayed as

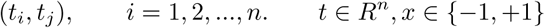

where *t*_*i*_ alludes to class marks and *i* signifies the quantity of information. The fundamental thought of this technique is to build the hyperplane that isolated the two classes of information with the condition

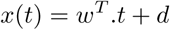

where *w* is perpendicular vector (n-dimensional) on the hyperplane and *d* is the bias. The expand of edge resembles limiting the heap standard ‖ *w* ‖^2^ characterized to

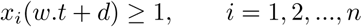

Using positive Lagrange multipliers *β*_*i*_ [13], i=1,2,…,n. The primal Lagrange function defined using above constraints as

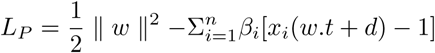

which is a convex quadratic programming problem to resolve the “dual” problem. Its means maximize *L*_*p*_ subject to the ∇*L*_*p*_ with significance *w* and *d* disappears. The solution of dual problem’s [13]

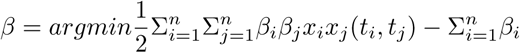

is given by above equation with constraints

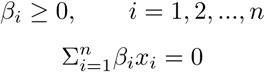

At that point, the nonzero *β*_*i*_ called support vectors, which are basic examples for grouping [17]. we’ll plan the training data to higher dimensional space to make a nonlinear choice surface. The isolating hyperplane to boost the edge and hyperplanes is spoken to kernel function as follows

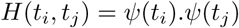

The decision function has characterized these kernels in the equation as [18]

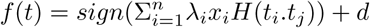

where *λ*_*i*_ are coefficients of Lagrange multipliers.

## III. Restricted Boltzmann

Restricted Boltzmann Machine (RBM) [19] [20] could be a two-layer undirected graphical model inside which the primary layer comprises of visible input variables **u** ∈ {0, 1}^*M*^, and a subsequent layer comprises of hidden variables (units) **v** ∈ {0, 1}^*N*^. We license only the between layer affiliations, i.e., there isn’t any relationship inside layers. Additionally, we include a third layer that speaks to recognizable yield variable *t* ∈ {1, 2, …, *R*}. we utilize the 1-to-R coding plan which at long last winds up in speaking to the yield as a twofold vector of length R meant by t, determined in the event that the yield (or class) is r, at that point all components are zero with the exception of component *t*_*r*_ which takes the worth 1.

A RBM with N hidden units might be a parametric model of the joint distribution of visible and hidden variables, that takes the structure [19]:

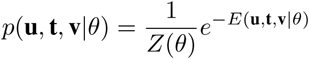

with parameter *θ* = {**b, c, d, W**^1^, **W**^2^}, and where:

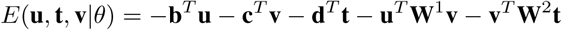

is the energy function, and

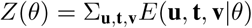

is a partition function. This model is termed Restricted Boltzmann Machine (RBM) [21]. The condition probability of a configuration o f t he v isible u nit, g iven a c onfigure of the hidden units **v**, is

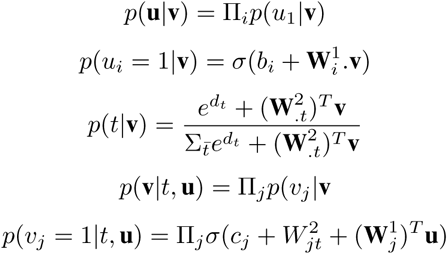

where *σ* is that the logistic sigmoid function, 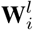 is that the ith row of weight matrix **W**^*l*^, 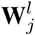 is that the jth column of weight matrix **W**^*l*^, 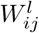 is the member of weights matrix **W**^*l*^. It is conceivable to precisely register the distribution *p*(*t*|**u**) which can be additionally acclimated pick the first likely class mark. This conditional distribution takes the accompanying structure

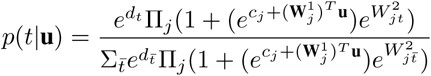

Pre-registering the terms 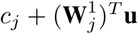 permits to slash back the time required for figuring the conditional distribution to *O*(*MD* + *MR*). inside the clinical setting, this possibility communicates the likelihood of happening ith contribution for the given different sources of info and class names.

Regularly, the boundary *θ* in RBM are gained from the information utilizing the probability work:

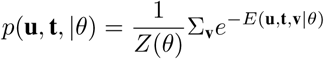

There are diverse inductive standards for learning RBM [19], [22]. The by and large used method is Contrastive Divergence [19], which is used here.

## IV. Deep belief networks (DBN)

DBN is balanced as neural frameworks, DBN has various non-straight covered layers, DBN is generatively pre-arranged near it can go about as non-direct dimensionality decline for input features vector, and over the long run, the organized teacher is another substantial information [23]. Let the yield neurons state y speak to the preparation model *y*^∗^. The inconsistency between the normal yield *y*^∗^ and accordingly the genuine yield *y* is utilizing the squared error measure:

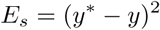

The adjustment in weight, which is added to the old weight, is satisfactory the product of the preparation rate and furthermore the gradient of the error function, increased by −1:

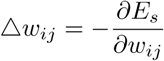

where the most of the dataset are unlabeled. Be that as it may, backpropagation neural system requires a labeled training data [24], [25]. As of late, consideration has moved to Deep learning [26] [27] [28] [29]. From Hinton’s point of view, the DBN is seen as an organization of simple learning modules every one of which can be a confined very RBM that contains a layer of obvious units. This layer speaks to information. Another layer of disguised units addresses includes that get higher-demand associations inside the data. the 2 layers are associated by a framework of evenly weighted associations (W) and there wear ‘t appear to be any associations inside a layer [30]. The key thought behind DBN is its weight (w), learned by an RBM define b oth *p*(*u*|*t, w*) then the prior distribution over hidden vectors *p*(*t*|*w*) [30]. The probability of generating a visual vector is written as

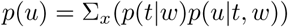

As the learning of DBN is likewise a computationally raised task, Hinton showed that RBMs could even be stacked and arranged in an exceedingly avaricious manner to make the DBN [31]. He presented a brisk calculation for learning DBN [31]. The heap update between visible u and hidden t joins essentially as

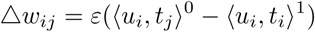

where 0 and 1 inside the above condition assign to the system information and reproduction state, separately.

## V. Experimental Results

We used Python 3.7.4 and import various libraries for data handling so that it can read datasets, then we’ve imported other libraries for data illustrate so that we are able to easily understand the data sets and interpret furthermore. We’ve identified NaN values throughout the row likewise as a column so as that we are able to fill it with various methods sort of a global mean or with the mean of row or column etc [32]. For organization, interpretation, presentation, and an intensive understanding of the information. We’ve described the knowledge set, so we could know the mean, variance, minimum, maximum limit, and thus, the precise distribution of knowledge set [33]. As we all know that ultimately our data is assessed into a binary classification which we’ve got illustrated the classification into a bar chart.

**Fig.1:**
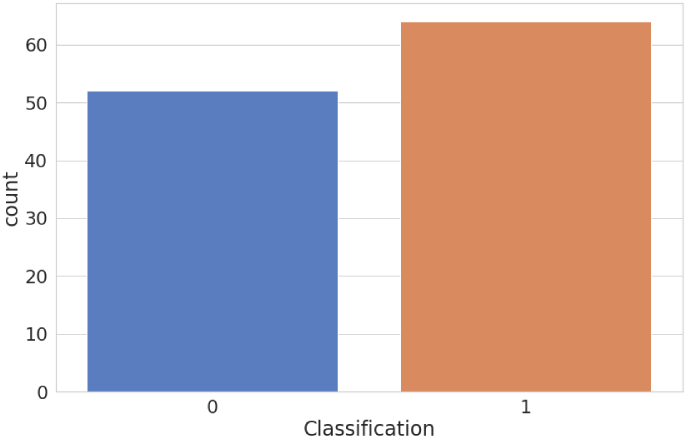
Classification labels

Descriptive statistics were calculated for all variables employed in the study using python syntax. we’ve got identified the assorted arrangement with relation to binary classification, which creates us know the mean, median, mode, and variance of a specific classification. As we respond to through observation, each data carries with it nine entry, and to grasp the distribution of feature in two classes, we’ve plated two or more plots in one figure so, that we are able to easily visualize each feature.

**Fig.2:**
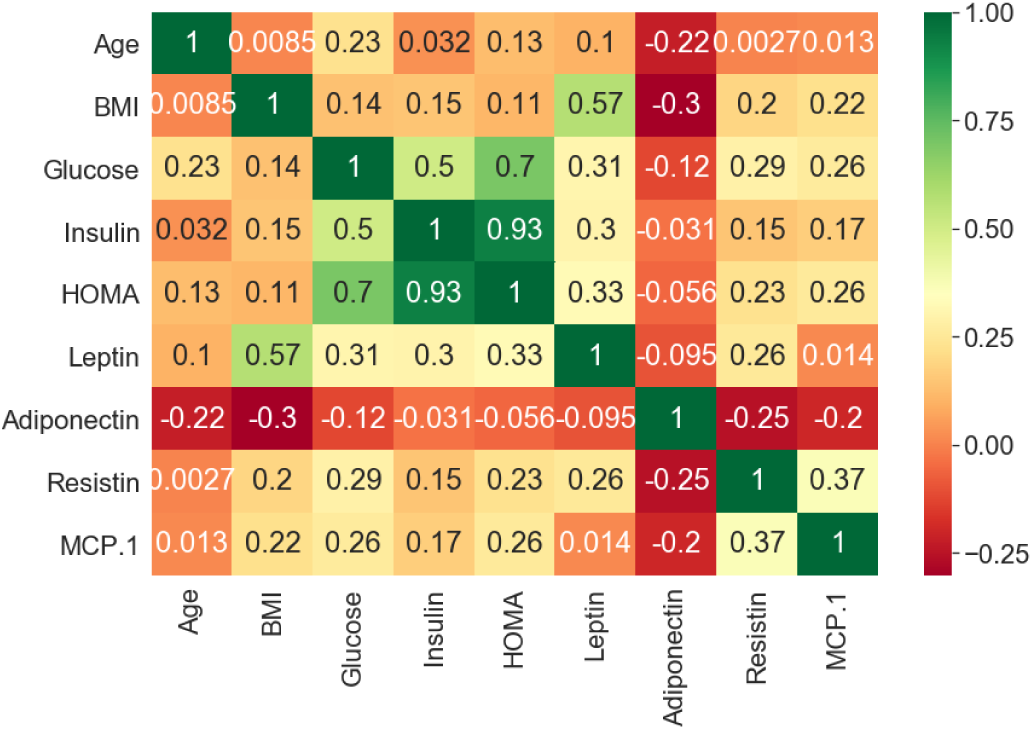
Heat Map/Correlation plot

Heat Map: We’ve got also used another technique to simplify the complex data set. A heat map [34] could be a two-dimensional portrayal of information inside which esteems are spoken to by color. There are regularly a few different ways to show heat maps, yet every one of them shares one thing in like manner. They use color to talk connections between information esteems that will be a lot harder to know whether introduced numerically in a surpassing spreadsheet. Here, we check how each feature vector is strongly correlated among them.

### A. Machine learning (ML) algorithms

We have used several supervised and unsupervised ML techniques to search out of the possible biomarker for breast cancer. to create a model for implementing the machine learning technique, Firstly we imported the library for the machine learning technique then we segregate data into tests and train to create a model. during this case, we’ve selected 25% data for testing and rest others for training.

Following Machine Learning techniques we’ve used with the subsequent accuracy:

**Table: 1.**

| Machine Learning Technique | Accuracy |
| --- | --- |
| Genetic Alogrithm | 0.875000 |
| Logistic Regression | 0.862069 |
| Linear SVM | 0.827586 |
| Random Forest | 0.793103 |
| Deep Belief Network | 0.758621 |
| Decision Tree | 0.724138 |
| Naive Bayes | 0.724138 |
| Radial SVM | 0.551724 |
| KNN | 0.551724 |

As we see the above Machine learning technique with their accuracy we found the Fluctuation within the accuracy of models is because of lack of knowledge. we’ve only 116 individuals data and to organize a decent model we want to data a minimum of in a thousand. The experimental data on cancer is extremely scarce. There are many various methods are available for solving these problems, but lack of knowledge is clearly not an answer. a technique to beat this problem is to either we’ve to rearrange the important data.

### B. Cross-validation

To overcome the matter with splitting into training and testing data sets, we’ve got applied K-fold cross-validation [35].To utilize an AI method, we’ve to indicate what division of information is utilized for training and testing informational collections and it’d be conceivable that some data in training data sets to urge the simplest results and a maximum number of knowledge points tested to urge the simplest validation [18]. Every information we’ve expelled from the training set into the test is lost for the training set. So, we’ve got to reset this swap. This can be the place cross-validation comes into the picture since this reason augments both of the sets, we have a decent arrangement of data inside the training sets to urge the most effective learning results, and also the maximum number of knowledge point we tested to induce to urge the simplest validation. The key thought of cross-validation is that segment of the information set into k canister of equivalent size, for instance: on the off chance that we have 300 datums and 10 container size which implies per receptacle have 30 information focuses, in this way, we’ve 30 information point in each container. So in cross-validation, one among the k subsets, as a testing test and remaining k-1 receptacle, are assembled as training sets at that point train AI calculations, and test the presentation on the testing set. [36]. AI calculations run various times, during this case, multiple times thus the normal the ten diverse testing sets exhibitions for the ten unique sets and normal the test results for those k experiments. this can be the way by which we will use all data sets for training, and everyone data for testing.

**Table: 2.**

| Machine Learning Technique | Cross-validation mean |
| --- | --- |
| Decision Tree | 0.581818 |
| Random Forest | 0.562879 |
| Naive Bayes | 0.547727 |
| Linear SVM | 0.533333 |
| Logistic Regression | 0.509091 |
| Radial SVM | 0.509091 |
| KNN | 0.506818 |

**Fig.3:**
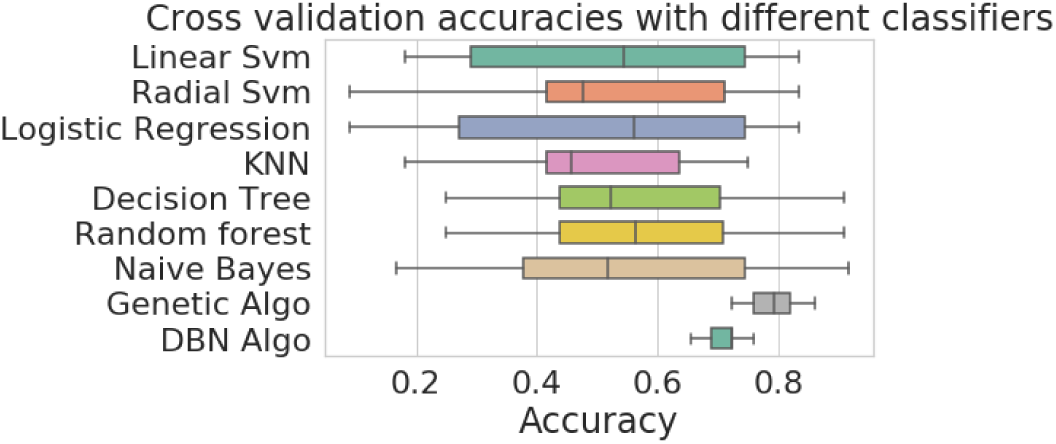
Cross Validation accuracies with different classifiers

### C. t-SNE Visualization

As we all know that we board a 3-D world in order that we are able to easily visualize the pattern within the 3-D, but what if we’ve data that is more complex? The visualization of knowledge is crucial steps to grasp the info and to grasp the complex data, dimension reduction is very important to step to require. Nowadays high dimensional data is everywhere and t-SNE [37] may be a popular technique for dimension reduction that we will use via sci-kit learn.

**Fig.4:**
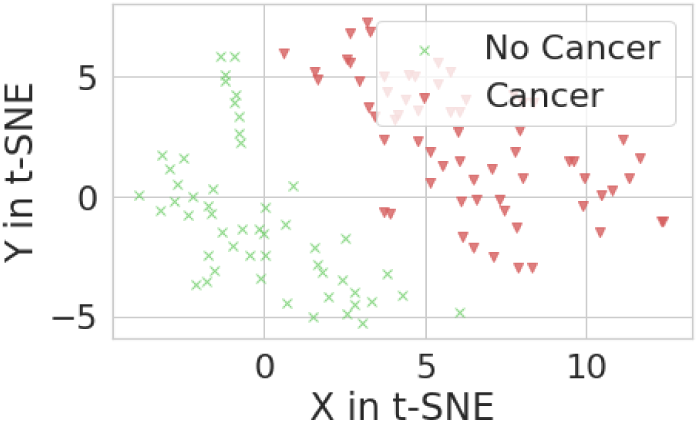
t-SNE visualization of Cancer data

The complete list of parameters which are important in predicting breast cancer are as follows

**Table: 3.**

| Feature | Importance | Normalized | Cumulative |
| --- | --- | --- | --- |
| Resistin | 33.4 | 0.213965 | 0.213965 |
| Age | 32.7 | 0.209481 | 0.423447 |
| Glucose | 31.9 | 0.204356 | 0.627803 |
| BMI | 16.9 | 0.108264 | 0.736067 |
| HOMA | 12.9 | 0.082639 | 0.818706 |
| Leptin | 10.5 | 0.067265 | 0.885971 |
| Adiponectin | 9.1 | 0.058296 | 0.944266 |
| Insulin | 4.5 | 0.028828 | 0.973094 |
| MCP.1 | 4.2 | 0.026906 | 1.000000 |

The feature analysis shows that there are not many highlights with the increasingly prescient incentives for the determination. From our investigation from Table:3, a few highlights like Resistin, Age, Glucose, and BMI are biomarkers for bosom malignant growth. Also, Patricio in his paper [7] discussed these features as the best biomarkers for prediction.

### D. ROC curve

We have tested a number of models to predict the possible outcomes and to understand what is the simplest model overall without having any quite hypothesis regarding the acceptance of the speed of the model. one all told the foremost credit risk models relies upon sensitivity and specificity [38] which has evaluated by the receiver operating graph i.e. ROC curve [39]. In this, a graph is plotted by plotting the sensitivity against specificity for every cutoff, the plot always starts with the lower-left corner where sensitivity is 0 and specificity is 1, in upper right corner correspond cut-off up to zero where sensitivity is 1 and specificity is 0. In general, speaking the maximum area of the best-left corner cover is healthier i.e. those curve cover maximum area of the best left have the next specificity associated with higher sensitivity [38]. The askew line partitioned the ROC curve into equal parts, a degree over the corner to corner for example upper left space represents good classification results contrariwise. Following formula for verity positive and false negative of ROC curve.

- 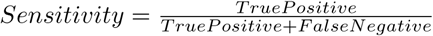
- 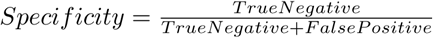

**Fig.5:**
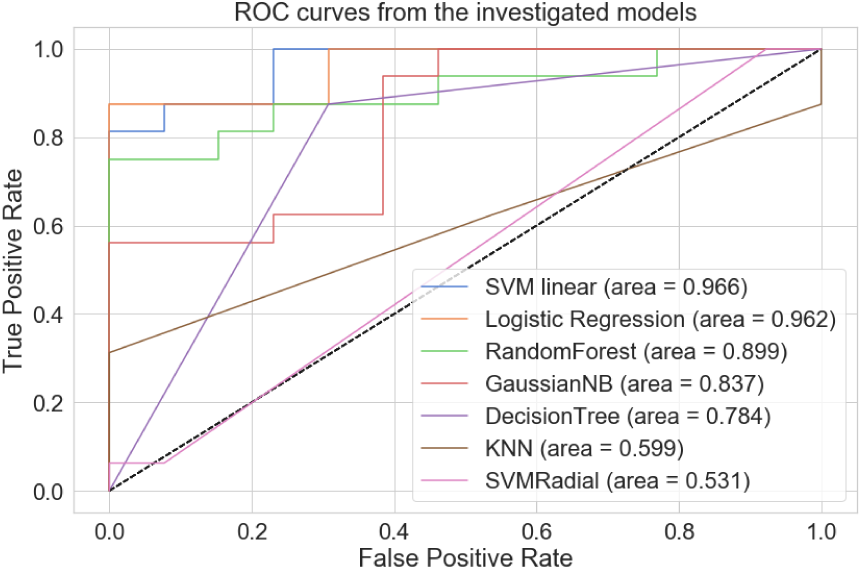
ROC curves from the investigated models

## VI. Conclusion

In this exploration, we’ve executed various strategies of a Machine learning tools for detecting breast cancer like deep belief nets, regression, decision tree, and genetic algorithms.. The overall neural network accuracy is 75.85% for DBN in breast cancer cases. Genetic algorithms give the upper accuracy of 87.50%. From Fig.3: We can observe that among all the classifiers, Deep Belief Nets and Genetic Algorithms are better classifiers then others. The variability is less in these two classifiers.Since the info sets utilized during this work are little if we have an oversized amount of information sets then methods used for prediction may be more accurate. We accept that the proposed framework is useful to the doctors for their official decisions on their patients. By utilizing such a technique, they’ll settle on exact decisions.

## Acknowledgment

KK thanks Department of Biotechnology (DBT),India: Joint DBT-Heidelberg University Graduate program on Big Data Research (fund: 2302086, assignment: 7815564) for support and RR is supported by the DST (Govt of India) through the JC Bose fellowship.

